# Topoisomerases I and II facilitate condensin DC translocation to organize and repress X chromosomes in *C. elegans*

**DOI:** 10.1101/2021.11.30.470639

**Authors:** Ana Karina Morao, Jun Kim, Daniel Obaji, Siyu Sun, Sevinc Ercan

## Abstract

Condensin complexes are evolutionarily conserved molecular motors that translocate along DNA and form loops. While condensin-mediated DNA looping is thought to direct the chain-passing activity of topoisomerase II to separate sister chromatids, it is not known if topological constraints in turn regulate loop formation *in vivo*. Here we applied auxin inducible degradation of topoisomerases I and II to determine how DNA topology affects the translocation of an X chromosome specific condensin that represses transcription for dosage compensation in *C. elegans* (condensin DC). We found that both topoisomerases colocalize with condensin DC and control its movement at different genomic scales. TOP-2 depletion hindered condensin DC translocation over long distances, resulting in accumulation around its X-specific recruitment sites and shorter Hi-C interactions. In contrast, TOP-1 depletion did not affect long-range spreading but resulted in accumulation of condensin DC within expressed gene bodies. Both TOP-1 and TOP-2 depletions resulted in X chromosome transcriptional upregulation indicating that condensin DC translocation at both scales is required for its function in gene repression. Together the distinct effects of TOP-1 and TOP-2 on condensin DC distribution revealed two distinct modes of condensin DC association with chromatin: long-range translocation that requires decatenation/unknotting of DNA and short-range translocation across genes that requires resolution of transcription-induced supercoiling.

## Introduction

Eukaryotic genome compaction is dynamically regulated throughout the cell cycle to accommodate essential genomic functions including gene expression during interphase and chromosome segregation during cell division. A key mechanism that achieves high levels of genome compaction is the formation of DNA loops. In addition to genome packaging, DNA looping modulates gene expression by bringing together regulatory sequences that are far apart in linear distance and by organizing the genome into functional domains that impact gene regulation (Merkenschlager and Nora, 2016). DNA looping is performed by an evolutionary conserved family of ring-shape motor proteins known as structural maintenance of chromosomes (SMC) complexes, which have been proposed to pull DNA through their opening in a process named “loop extrusion” (Uhlmann, 2016; Banigan and Mirny, 2020). Recently, this model was supported by single molecule experiments allowing the direct imaging of DNA loop extrusion by the SMC complexes condensin and cohesin *in vitro* (Ganji et al., 2018; Golfier et al., 2020; Kong et al., 2020).

We reason that *in vivo* SMC complexes may be challenged by DNA supercoiling and entanglements arising from transcription and replication. Such DNA templated processes involve opening of the double helix and translocation of large protein complexes that force DNA to rotate around its axis. This unwinding and rotation generate negative and positive supercoiling behind and ahead the translocating machineries, respectively (Liu and Wang, 1987). In addition to supercoiling, DNA molecules can be entangled especially after replication where positive supercoiling at the replication fork causes the intertwining of the newly replicated duplexes leading to their eventual catenation (Nitiss, 2009). Topological stress is resolved by topoisomerases, a family of enzymes with DNA cleavage and re-ligation activity. Topoisomerases are classified based on whether they cleave one or two DNA strands in type I (e.g. topoisomerase I) and type II (e.g. topoisomerase II) (Baranello et al., 2013). While both topoisomerase I and II can resolve positive and negative supercoils, TOP-1 has been described to have a major role in the relaxation of transcription-induced supercoiling (Durand-Dubief et al., 2010; Teves and Henikoff, 2014). Unlike TOP-1, TOP-2 is able to resolve DNA catenations that require its double-stranded DNA cutting and passage activity (Baranello et al., 2013).

Inactivation of topoisomerase II and condensin leads to two main phenotypes: impaired chromosome compaction and failure to resolve sister chromatids, causing chromosome bridges and segregation defects (Uhlmann, 2016; Piskadlo and Oliveira, 2017). Experimental data in different organisms showed accumulation of topoisomerase II induced DNA entanglements in the absence of condensin, thus suggesting that condensin is required to direct the activity of topoisomerase II towards the disentanglement of DNA molecules (Baxter et al., 2011; Charbin et al., 2014; Dyson et al., 2020; Piskadlo et al., 2017; Sen et al., 2016). Computational simulations suggest that the mechanism underlying the cooperative activity of SMC complexes and topoisomerase II stems from the interplay between their respective loop extrusion and DNA-duplex passage activities (Goloborodko et al., 2016). Subsequently, it was proposed that topological entanglements resulting from DNA transactions such as transcription and replication, would get pushed and spatially localized by the process of loop extrusion, and that this would promote chromatin unknotting by topoisomerase II (Orlandini et al., 2019; Racko et al., 2018). Experimental support for this model came from the observation that mammalian topoisomerase II beta binds to CTCF/cohesin occupied regions in different cell lines (Canela et al., 2017; Manville et al., 2015; Uusküla-Reimand et al., 2016) and induces DSBs at loop anchors (Canela et al., 2019, 2017; Gothe et al., 2019). While the role of condensin in directing topoisomerase II to decatenate sister chromatids is better understood, how DNA entanglements and supercoiling affect condensin-mediated chromosome compaction *in vivo* remains unknown.

To address how condensin translocation is modulated by DNA topology *in vivo*, a system that allows measurement of the direction and the extent of condensin translocation is needed. Such a unique eukaryotic system exists for a specialized *C. elegans* condensin that forms the core of the X chromosome <u>**D**</u>osage <u>**C**</u>ompensation complex (Ercan, 2015; Meyer, 2005) (**Figure 1A**). This condensin, named condensin DC, is exclusively recruited to the two X chromosomes in hermaphrodites by the proteins SDC-2, SDC-3 and DPY-30, which are enriched at a small number of sequence specific <u>**r**</u>ecruitment <u>**e**</u>lements on the <u>**X**</u> (*rex* sites) (Albritton et al., 2017; Ercan et al., 2007; Jans et al., 2009; McDonel et al., 2006; Meyer, 2010). Condensin DC spreads linearly along the X chromosome and concentrates at active promoters and other accessible gene regulatory elements (Ercan et al., 2009; Jans et al., 2009; Street et al., 2019). The action of condensin DC halves transcription initiation of both X chromosomes in hermaphrodites to equalize it to the single X present in males (Kramer et al., 2015; Kruesi et al., 2013) and remodels their 3D structure. (Anderson et al., 2019; Crane et al., 2015; Jimenez et al., 2021). The X-specific recruitment and linear spreading of condensin DC, along with the presence of the strong *rex* sites acting as TAD boundaries allows the analysis of condensin DC translocation and loop formation by ChIP-seq and Hi-C, respectively.

**Figure 1.**
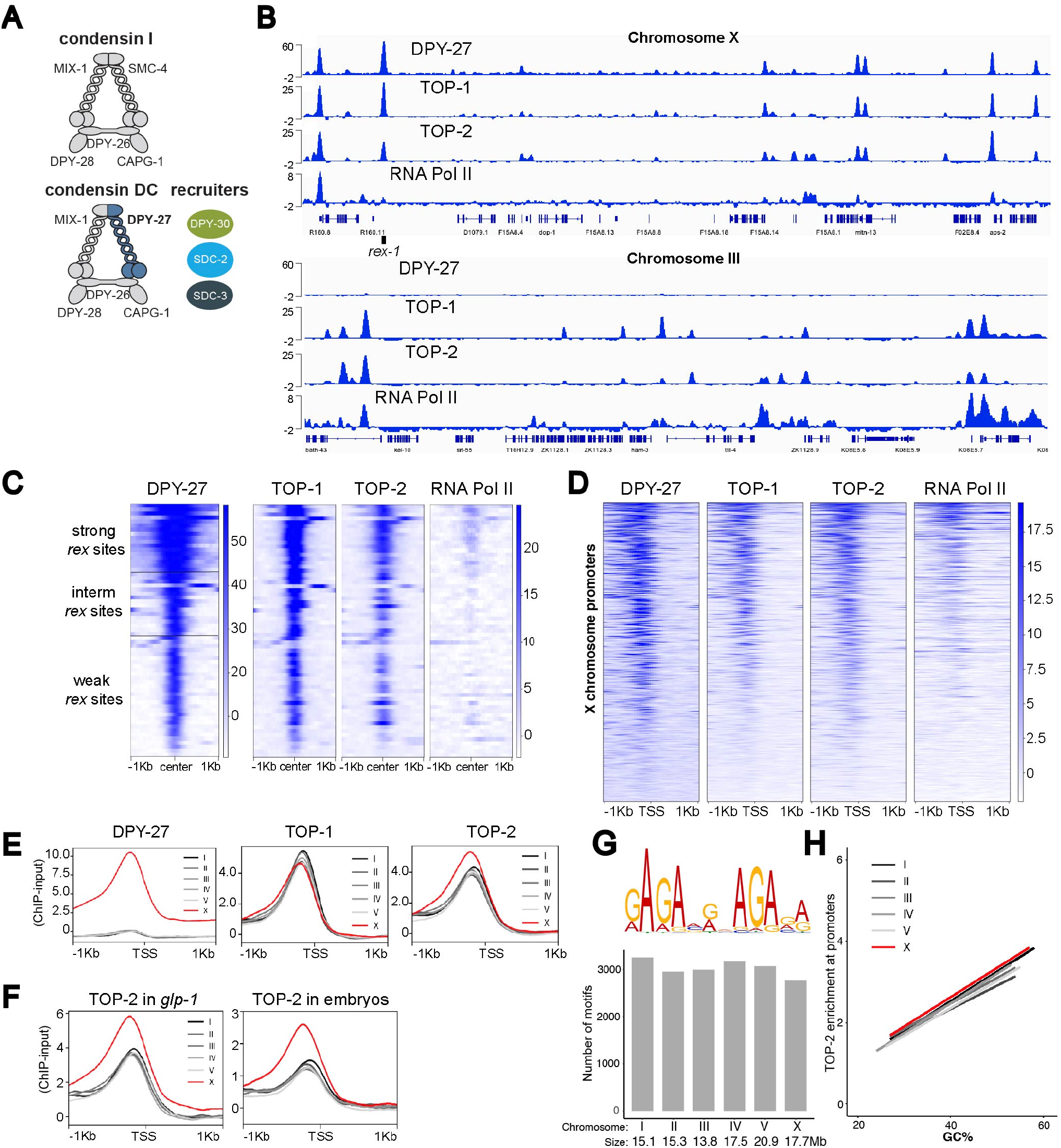
TOP-1 and TOP-2 overlap with condensin DC on the X chromosome and bind to active promoters across the genome. A) Schematic representation of the *C. elegans* condensin I and condensin DC complexes. Condensin DC differs from condensin I by one subunit, the SMC-4 variant, DPY-27. Binding of condensin DC to the X chromosomes requires the recruiter proteins: SDC-2, SDC-3 and DPY-30. B) DPY-27, TOP-1, TOP-2 and RNA Pol-II ChIP-seq profiles at representative regions on chromosomes X (top) and III (bottom). Normalized ChIP-input coverage is shown. Genes are displayed and the location of *rex-1* is indicated. C) Heatmap showing ChIP-seq signal for DPY-27, TOP-1, TOP-2 and RNA Pol-II across a 2 Kb window centered around 64 *rex* sites described in Albritton et al., 2017. *rex* sites are classified into three strength categories (weak, intermediate and strong) based on ChIP-seq enrichment of condensin DC subunits and recruiter proteins (Albritton et al., 2017). TOP-1 and TOP-2 binding intensity correlates with condensin DC. RNA Pol-II is mostly absent from *rex* sites. D) Heatmap showing DPY-27, TOP-1, TOP-2 and RNA Pol-II ChIP-seq signals across X chromosome transcription start sites (TSS) defined by Kruesi et al., 2013 using GRO-seq. DPY-27, TOP-1 and TOP-2 binding correlate with RNA Pol-II. E) Average DPY-27, TOP-1 and TOP-2 ChIP-seq scores are plotted across a 2 Kb window centered around TSSs on X and autosomes. TOP-1 and TOP-2 bind promoters across all chromosomes but TOP-2 shows higher binding on X chromosome promoters. F) Average TOP-2 ChIP-seq scores are plotted across TSSs on X and autosomes in *glp-1(q224)* adults lacking a germline and WT embryos. TOP-2 X enrichment is observed in both conditions. G) Top: MEME (version 5.3.3) was used to perform a motif search across 150 bp regions centered on TOP-2 ChIP-seq summits in L3. The most significant DNA motif found is shown. Bottom: The *C. elegans* genome (ce10) was scanned for instances of the top DNA motif using PWMScan. The number of hits (p-value cutoff 1e-5) obtained per chromosome are shown. The TOP-2 DNA motif is not enriched on the X chromosome. H) The GC content of 250 bp windows centered on TSSs was plotted on the x axis and TOP-2 ChIP-seq scores across the same window were plotted on the Y axis. Data was fitted to a linear model. TOP-2 binding positively correlates with GC content. For a given GC value, TOP-2 ChIP-seq scores are higher on the X compared to autosomes.

Here we analyzed condensin DC binding and function in regulating 3D contacts and transcription, upon auxin-inducible depletion of TOP-1 and TOP-2. We found that topoisomerases I and II colocalize with condensin DC on chromatin and are required for its translocation at different scales. TOP-2 depletion hindered the long-range spreading of condensin DC, resulting in increased condensin DC binding around the strong *rex* sites. Reduced condensin DC spreading resulted in shorter Hi-C contacts on the X chromosome. TOP-1 depletion resulted in gene-body accumulation of condensin DC and RNA Pol II. X chromosomes were derepressed in both TOP-1 and TOP-2 depleted conditions, thus condensin DC mediated transcriptional repression involves both long-range and local translocation. Importantly, TOP-1 depletion did not reduce the range of condensin DC spreading from the strong *rex* sites, revealing two distinct modes of condensin DC translocation that are sensitive to different types of topological constraints. Based on these results, we propose that the unknotting activity of TOP-2 is required for the translocation of condensin DC across long distances, whereas local translocation across gene bodies requires the resolution of transcription-induced supercoiling performed mainly by TOP-1.

## Results

### TOP-1 and TOP-2 localize to active promoters across the genome and overlap with condensin DC on the X

We first reasoned that if topoisomerases regulate condensin DC translocation on the X chromosomes, they may co-localize on chromatin. To test this, we determined the binding profile of the two major somatic topoisomerases in *C. elegans*, TOP-1 (M01E5.5) and TOP-2 (K12D12.1), in L2/L3 worms by chromatin immunoprecipitation followed by sequencing (ChIP-seq) (Jaramillo-Lambert et al., 2016; Lee et al., 1998). For TOP-2, we used a published strain, in which the endogenous *top-2* gene is tagged with GFP (Ladouceur et al., 2017). *C. elegans* TOP-1 has two isoforms, TOP-1α, which contains five exons and TOP-1β, which lacks the second exon (Lee et al., 1998). To tag both isoforms, we introduced a degron-GFP tag at the end of exon five using CRISPR-Cas9.

The ChIP-seq profile of DPY-27, the condensin DC SMC-4 variant, shows strong enrichment at recruitment sites (*rex*), different levels of moderate enrichment at promoters and uniform baseline signal across the X chromosome (Ercan et al 2007, Jans et al, 2009, Street et al, 2019) (**Figure 1B**). In line with a functional interplay between condensin DC and topoisomerases, both TOP-1 and TOP-2 ChIP-seq signal followed that of the condensin DC (**Figure 1B** top panel). TOP-1 and TOP-2 were enriched at *rex* sites (**Figure 1C**), and bound to X chromosome promoters, in a manner largely proportional to condensin DC binding (**Figure 1D**). Thus, topoisomerases I and II co-localize with condensin DC on the X chromosome.

Unlike condensin DC, TOP-1 and TOP-2 also bind to the autosomal promoters (**Figure 1B** low panel). As previously described in other organisms (Durand-Dubief et al., 2010; Heldrich et al., 2020; Uusküla-Reimand et al., 2016), TOP-1 and TOP-2 binding in *C. elegans* correlated with transcriptional activity (**Figure 1D**).

### TOP-2 binding is enriched at X chromosome promoters

Strikingly, analysis of topoisomerase binding across promoters showed stronger TOP-2 binding on the X compared to autosomes (**Figure 1E and S1A**). Unlike TOP-2, TOP-1 was not enriched on the X (**Figure 1E**). TOP-2 and condensin have been historically identified as key players in the resolution and compaction of mitotic chromosomes. Here, we also see a specific association between TOP-2 and an X-specific condensin, highlighting the relevancy of our system for analyzing the function of TOP-2 and condensins in chromosome compaction.

In *C. elegans*, condensin DC is only expressed in somatic cells, therefore if X enrichment of TOP-2 is related to condensin DC function, it should happen in the soma (Strome et al., 2014). Indeed, TOP-2 ChIP-seq showed X enrichment in *glp-1* mutant adults (**Figure 1F**), which lack the germline when grown at the restrictive temperature (**Figure S1B**). X-enrichment of TOP-2 binding was also observed in mixed-stage embryos containing only two germ cells, suggesting that like condensin DC, TOP-2 is enriched on X chromosomes in somatic cells throughout development (**Figure 1F**).

To address if there are X-specific sequence features that recruit TOP-2, we performed a motif search under TOP-2 binding peaks across the genome. This analysis yielded one prominent motif (**Figure 1G**). We then scanned the genome for the motif sequence and found that it was not enriched on the X chromosome (**Figure 1G**). The next sequence feature we considered is GC content, as *C. elegans* X chromosome promoters have higher GC content than autosomes (Ercan et al., 2011). While TOP-2 binding across all chromosomes correlates positively with GC content (**Figure 1H)**, for sites with the same GC content, X chromosomal binding was higher than that of autosomes. Thus, increased GC content of X chromosomal promoters does not underlie TOP-2 enrichment on the X (**Figure 1H)**. Together, our results support the conclusion that TOP-2 binding is increased on the dosage compensated X chromosomes bound by condensin DC.

### Recruitment elements on the X *(rex)* sites recruit TOP-2

We next hypothesized that condensin DC might be responsible for the X chromosome enrichment of TOP-2. In bacteria, the SMC complex MukBEF recruits topoisomerase IV to *ori* regions (Nicolas et al., 2014) and in mammalian cells, cohesin interacts with topoisomerase II (Uusküla-Reimand et al., 2016) and is required for TOP-2 binding over loop anchors (Canela et al., 2019). To address if condensin DC is required for TOP-2 binding on the X, we introduced a degron-GFP tag at the C-terminus of DPY-27 and used the Auxin Inducible Degradation system by expressing the TIR-1 component of the degradation machinery exclusively in somatic cells where condensin DC is present (Zhang et al., 2015) (**Figure 2A**). Insertion of the degron-GFP tag in the presence of TIR-1 already impaired DPY-27 function resulting in reduced binding and X chromosome transcriptional upregulation without the addition of auxin (**Figure S2A, S2B and S2C**). Auxin treatment further reduced DPY-27 protein amount, eliminating binding to the X chromosomes (**Figure 2B, S2A and S2B**). In both partial and complete DPY-27 depletion conditions, TOP-2 enrichment on the X chromosome promoters was maintained (**Figure 2C**).

**Figure 2.**
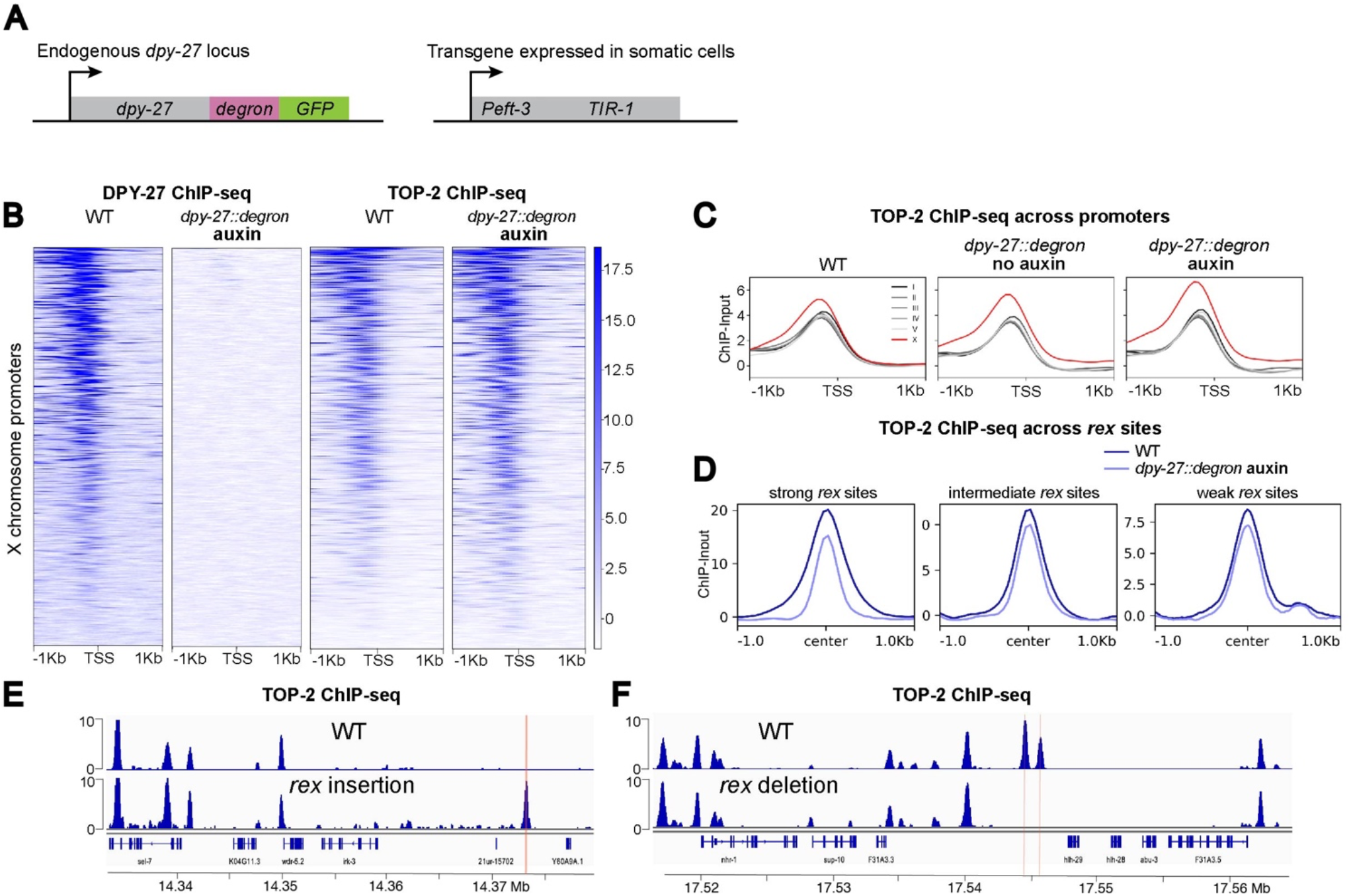
Recruitment elements on the X *(rex)* sites recruit TOP-2. A) Schematic representation of components of the Auxin Inducible Degradation system used for DPY-27 depletion. B) Heatmap showing DPY-27 and TOP-2 ChIP-seq signals around X chromosome TSSs in WT and DPY-27 depletion conditions. *dpy-27::degron* worms were treated with auxin for 60 min. In the absence of DPY-27, TOP-2 binding to X promoters is maintained. C) Average TOP-2 ChIP-seq scores in WT, DPY-27 partial (no-auxin) and complete (auxin) depletion conditions are plotted across a 2 Kb window centered around TSSs on X and autosomes. TOP-2 enrichment on X chromosome promoters is maintained in the absence of DPY-27. D) Average TOP-2 ChIP-seq scores in WT and DPY-27 depletion conditions are plotted across a 2 Kb window centered at strong, intermediate and weak *rex* sites. E) ChIP-seq profiles comparing TOP-2 signal in wild-type with a strain carrying an ectopic *rex-8* insertion on the X chromosome. Location of the insertion is indicated by a red line. F) ChIP-seq profiles comparing TOP-2 signal in wild-type with a strain in which ∼100 bp containing the two recruiting motifs of the endogenous *rex-41* have been deleted. Locations of the deletions are indicated by red lines.

Interestingly, despite the maintenance of TOP-2 binding at promoters, its binding over strong *rex* sites was markedly reduced in the absence of condensin DC, whereas enrichment at intermediate and weak *rex* sites was less affected (**Figure 2D**). Strong recruitment sites consist of interspaced clusters of a 12 bp motif that are important for condensin DC recruitment (Albritton et al., 2017; Ercan et al., 2007; Jans et al., 2009) and form the loop anchors at TAD boundaries on the X (Anderson et al., 2019; Crane et al., 2015; Jimenez et al., 2021; Rowley et al., 2020). To test if *rex* sites recruit TOP-2, we asked if insertion of a strong *rex* at an ectopic location on the X that normally lacks TOP-2 could create a new binding site (Albritton et al., 2017; Jimenez et al., 2021). Indeed, TOP-2 ChIP-seq showed a new peak at the insertion site of an extra copy of *rex-8* (358 bp), indicating that a strong *rex* is sufficient to recruit TOP-2 (**Figure 2E)**. IgG and RNA Pol-II ChIP-seq signal showed background levels at the insertion site indicating that TOP-2 recruitment is specific (**Figure S2D**). To determine if *rex* sequences are necessary for TOP-2 binding to the *rex* sites, we performed TOP-2 ChIP-seq in a strain in which ∼100 bp containing the two recruiting motifs of the endogenous *rex-41* have been deleted (Albritton et al., 2017). Here, the TOP-2 binding domain spanning around 2.5 Kb completely disappeared (**Figure 2F)**. Thus, *rex* sequences are sufficient and required for TOP2 binding to the *rex* sites.

It remains unclear however if it is condensin DC or another protein functioning at the *rex* sites that recruits TOP-2. One hypothesis is that condensin DC recruitment and spreading from the *rex* sites causes the elevated TOP-2 binding at the X chromosomes. It is possible that increased transcription of the X chromosomes upon DPY-27 depletion countered the potential reduction in TOP-2 binding (**Figure S2C)**. Indeed, TOP-2 binding is correlated with transcription, thus derepression of the X is expected to increase TOP-2 binding specifically at the promoters but not *rex* sites, which is what we observed. Thus, it is possible that condensin DC activity increases TOP-2 binding to the X chromosomes.

### TOP-1 and TOP-2 depletion results in X chromosome transcriptional upregulation

To determine the functional impact of topoisomerase I and II on condensin DC mediated chromosome-wide transcriptional repression, we performed mRNA-seq in TOP-1 and TOP-2 depleted conditions. For this, we endogenously tagged *top-1* and *top-2* with degron-GFP using CRISPR/Cas9 in the strain expressing TIR-1 in somatic cells (**Figure 3A**). To avoid affecting cell division, we performed short-term depletion and used L2/L3 larvae since most somatic cells are no longer dividing at this developmental stage (Sulston and Horvitz, 1977). For both TOP-1 and TOP-2, nuclear GFP signal could no longer be detected in somatic cells after one hour of auxin treatment (**Figure 3B**). We performed mRNA-seq after thirty minutes, one hour and two hours of auxin-mediated TOP-1 or TOP-2 degradation. As controls, we used *top-1* or *top-2* degron-tagged worms that were not treated with auxin (no-auxin samples) (**Figure 3A**), as well as a strain lacking the degron-GFP tag but containing the TIR-1 transgene, that was treated with auxin (no-tag auxin) (**Figure 3A**).

**Figure 3.**
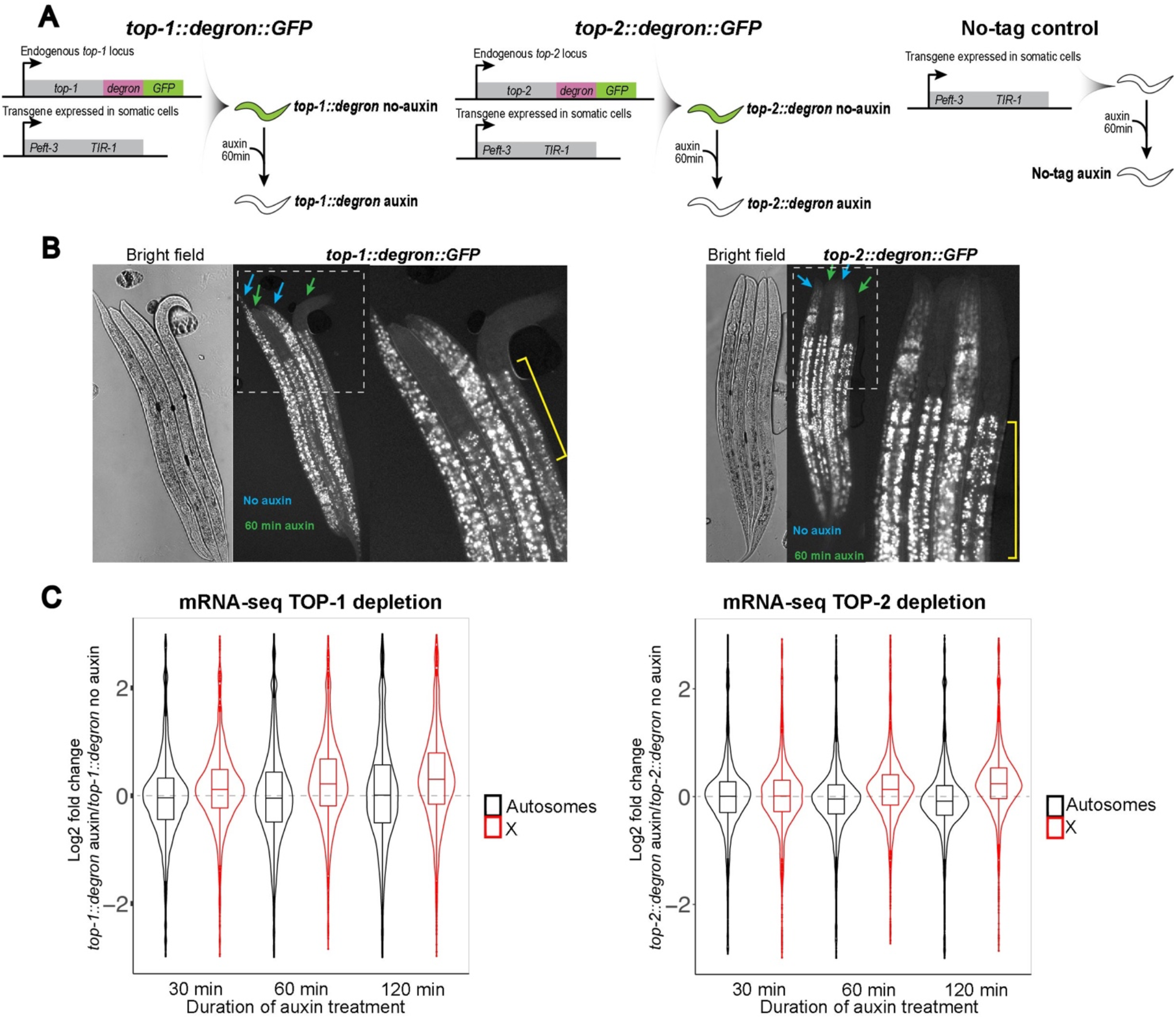
TOP-1 and TOP-2 depletion results in X chromosome transcriptional upregulation. A) Schematic representation of components of the Auxin Inducible Degradation system used for TOP-1 and TOP-2 depletion experiments. As controls, we used *top-1* or *top-2* degron-tagged worms that were not treated with auxin (no-auxin samples), as well as a strain lacking the degron-GFP tag but containing the TIR-1 transgene, that was treated with auxin (no-tag auxin). B) Auxin induced degradation of TOP-1 (left) and TOP-2 (right) in somatic cells. *top-1::degron::GFP* and *top-2::degron::GFP* L3 worms were incubated in auxin (green arrows) and no-auxin (blue arrows) plates for one hour, followed by imaging. Rightmost images show higher magnification views of the outline regions. Yellow brackets indicate autofluorescent gut granules (Hermann et al., 2005). C) mRNA-seq was performed after 30 min, 60 min and 120 min auxin-mediated depletion of TOP-1(left) and TOP-2 (right). Distribution of log2 fold changes between the auxin treated and no-auxin conditions are shown for autosomes and X chromosomes. Depletion of both TOP-1 and TOP-2 cause X chromosome derepression.

In both TOP-1 and TOP-2 depleted conditions, average X chromosomal gene expression increased progressively with longer auxin incubation time, while autosomes were less affected (**Figure 3C and S3)**. These results indicate that TOP-1 and TOP-2 are important for the repression of X chromosomes. While upregulation of the X chromosome upon TOP-2 depletion supports the idea that X-enrichment of TOP-2 is important for repression, the fact that TOP-1 also showed upregulation suggested a broader role for topoisomerases in X chromosome repression. Next, we explored the mechanisms of these roles.

### TOP-2 depletion reduces the size of condensin DC mediated DNA contacts on the X chromosomes

Loop extrusion performed by SMC complexes and double-stranded DNA passage performed by topoisomerase II have been proposed to contribute to the 3D genome organization throughout the cell cycle (Björkegren and Baranello, 2018; Piskadlo and Oliveira, 2017). Here, we reasoned that TOP-2 may regulate the processivity of loop extrusion by condensin DC through the resolution of chromatin catenations and knots. To test this idea, we performed Hi-C in larvae depleted of TOP-2 by one hour of auxin treatment. Upon TOP-2 knockdown, the TADs on the X appeared weaker, as seen by decreased insulation at TAD boundaries along the chromosome (**Figure 4A**). There was no strong effect on autosomes, indicating that TOP-2 is especially required for X chromosome 3D organization (**Figure 4A**).

**Figure 4.**
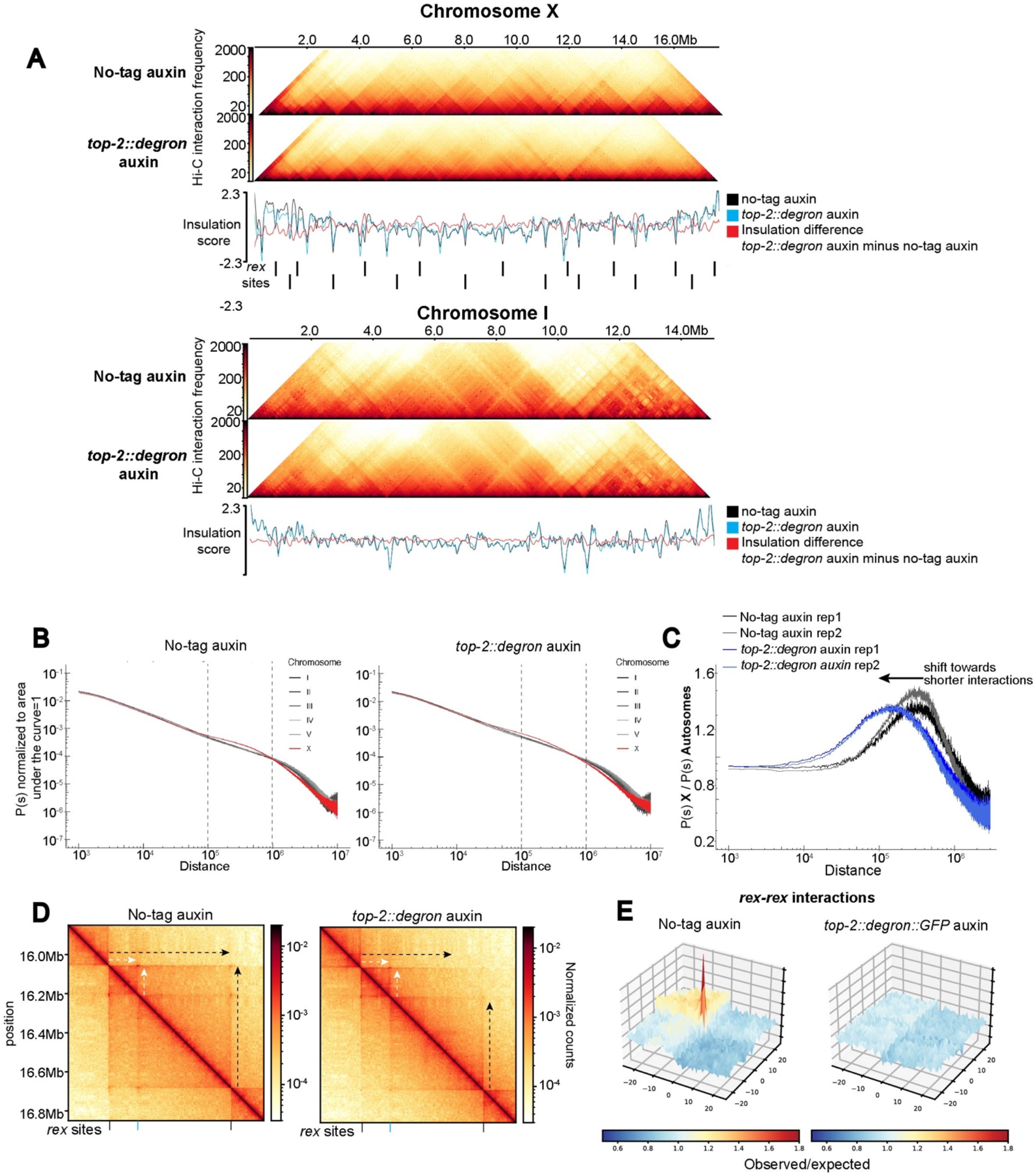
TOP-2 depletion leads to shorter DNA interactions on the X chromosomes. A) Hi-C heatmap and insulation scores of chromosomes X and I in no-tag auxin and *top-2::degron* auxin conditions. The positions of the 17 strong *rex* sites described in Albritton et al,. 2017 are indicated. The insulation scores and their subtractions are shown below. B) Distance decay curve showing the relationship between 500 bp binned genomic distance and average contact probability computed per chromosome in the no-tag auxin control and *top-2::degron* auxin. Dashed lines in both panels highlight the distance range of interactions that are enriched on the X chromosome in the control condition (100Kb-1Mb). C) X-enriched chromosomal contacts are visualized by autosome normalized distance decay curves. For every genomic distance, the unity normalized contact probability of the X-chromosome, P(s) X, is divided by that of the autosomes, P(s) Autosomes. UponTOP-2 depletion, interactions enriched on the X get shorter. D) Hi-C snapshot of 1MB region on X-chromosome. Black arrows indicate long stripes emanating from strong *rex* sites. White arrows indicate short stripes of nested TAD. Strong (black line) and intermediate (blue line) *rex* sites are indicated at the bottom. E) Meta-’dot’ plot showing the average strength of interactions between pairs of *rex* sites on a distance-normalized matrix. For 17 strong *rex* sites, a total of 33 *rex-rex* pairs located within 3 Mb of each other were used.

Within the frame of the loop extrusion hypothesis, if TOP-2 is required for condensin-DC processivity, condensin-DC mediated 3D DNA contacts on the X chromosome should shorten upon TOP-2 depletion. Indeed, Hi-C distance decay curves, where contact probabilities are plotted as a function of genomic distance, showed reduced distance-range of interactions on the X (**Figure 4B**). In control larvae, autosomes curves align together reflecting a similar range of interactions, while X displays increased interactions in the distance range of TADs (∼100 Kb - 1Mb) producing a characteristic hump shape that is not observed for the autosomes (**Figure 4B** left panel). TOP-2 degradation led to a leftward shift of the hump, indicating a shift towards shorter-range interactions (**Figure 4B** right panel). Autosomes also showed a shift toward shorter interactions but to a lesser extent than the X (**Figure S4A**). To isolate the condensin-DC specific effect of TOP-2 depletion, we divided the distance decay curve of the X chromosome by that of autosomes. In the control, normalization to autosomes highlighted X-specific enrichment of long-range 3D contacts centering approximately ∼300 Kb (**Figure 4C)**. In the TOP-2 depleted condition X-enriched 3D contacts shift towards shorter interactions centering ∼100 Kb (**Figure 4C**). These results indicate that condensin DC makes shorter loops in the absence of TOP-2, consistent with the hypothesis that removal of DNA catenations is important for the processivity of condensin DC loop extrusion.

A second prediction of reduced condensin DC processivity is that ‘stripes’ should get shorter upon TOP-2 depletion. Stripes are produced by the asymmetric reeling-in of DNA through the SMC ring, which results in a single locus (anchor) forming interactions with progressively more distant loci (Fudenberg et al., 2017; Vian et al., 2018). On the X, stripes are anchored at *rex* sites which correspond to TAD boundaries (Anderson et al., 2019; Crane et al., 2015; Jimenez et al., 2021). In line with hindered loop extrusion, stripes anchored at *rex* sites got shorter upon TOP-2 depletion. A clear example is shown in Figure 4D, where stripes of around 600 Kb originating from two strong *rex* sites are reduced (**Figure 4D**, black arrows), whereas stripes flanking a smaller nested TAD are not affected (**Figure 4D**, white arrows). A metaplot analysis of interactions around the 17 strong *rex* sites shows the overall shortening of stripes emanating from strong *rex* sites (**Figure S4B**). Since the strong *rex* sites are stripe anchors, they are brought together in 3D space. Indeed, *rex-rex* interactions are among the most frequent long-range interactions on the X chromosome (Anderson et al., 2019; Crane et al., 2015). Consistent with shorter stripes, we observe that upon TOP-2 depletion, *rex-rex* interactions were lost (**Figure 4E**). Thus, both predictions of the hypothesis, shorter X-specific 3D DNA contacts and shorter stripes, are met. Therefore, we conclude that TOP-2 is required for condensin DC mediated 3D organization of the X chromosomes by increasing the processivity of condensin DC translocation.

### TOP-2 degradation reduces the spreading of condensin DC along the X chromosome

To directly test the idea that TOP-2 is required for condensin DC processivity, we analyzed condensin DC binding by ChIP-seq upon one hour of TOP-2 depletion. We reasoned that in the absence of TOP-2, condensin DC binding should decline with increasing distance from the *rex* sites, as a result of hindered spreading. DPY-27 ChIP-seq in the no-tag auxin and no-auxin controls shows strong and narrow enrichment at *rex* sites and moderate enrichment that is evenly distributed across the X (**Figure 5A, Figure S5**). Upon TOP-2 degradation, DPY-27 binding became concentrated around the strong *rex* sites and progressively decreased in a distance dependent manner creating “mountains” of condensin DC across the entire X chromosome that were centered around strong *rex* sites (**Figure 5A**).

**Figure 5.**
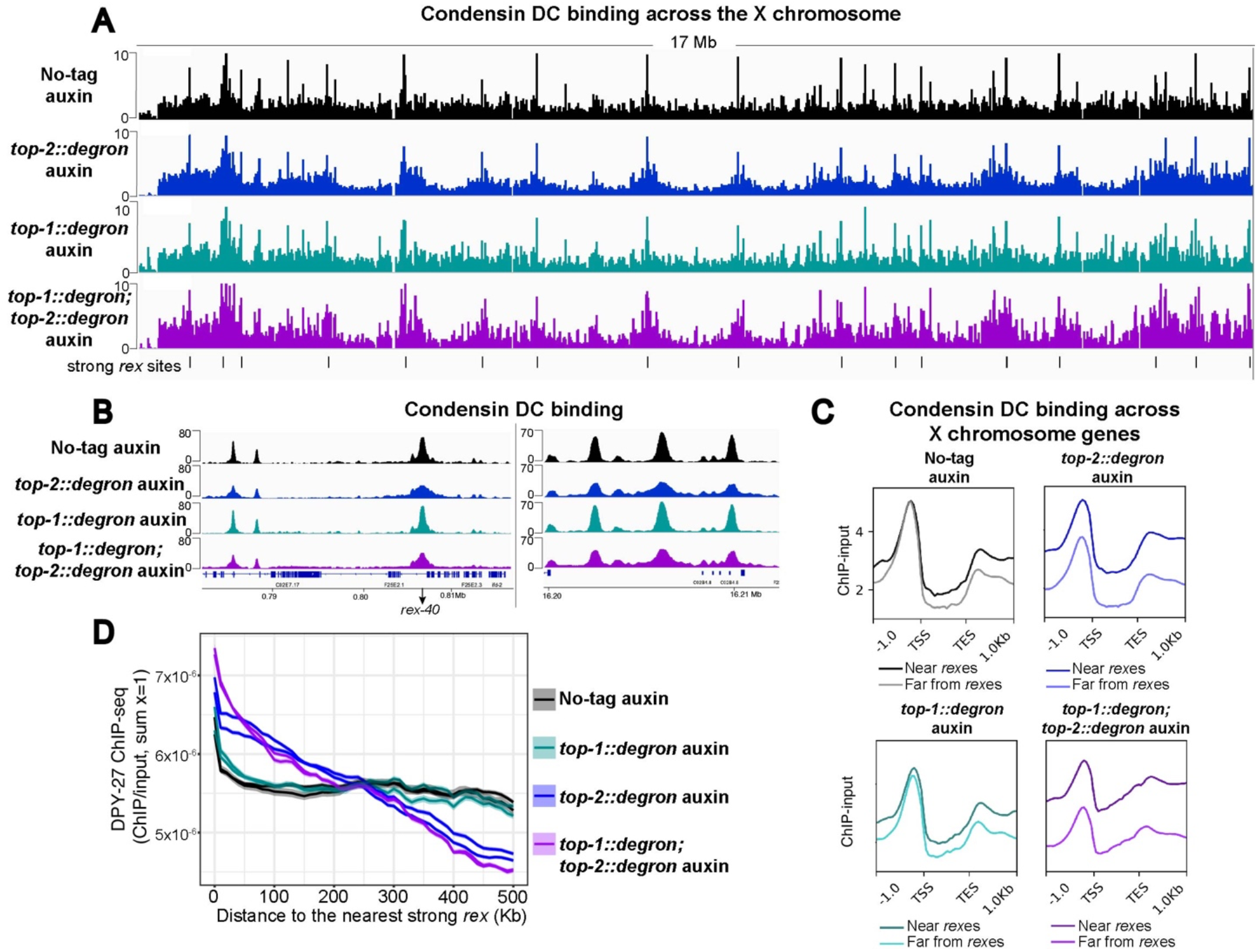
Spreading of condensin DC is hindered in the absence of TOP-2. A) X chromosome view of DPY-27 ChIP-seq profile in control (no-tag auxin), TOP-1, TOP-2 and double TOP-1/TOP-2 depletions. All the strains were treated with 1mM auxin for 60 minutes. Black lines at the bottom indicate the location of strong *rex* sites. Normalized ChIP-Input coverage is shown. B) DPY-27 ChIP-seq profile in control, TOP-1, TOP-2 and double TOP-1/TOP-2 depletions at representative regions of the X chromosome. C) Genes on the X chromosome were classified according to their distance to a strong *rex* site into two categories: near *rex* sites (within 400Kb of a strong *rex* site) and far from *rex* sites (more than 400Kb away from a strong *rex* site). Average DPY-27 ChIP-seq scores in control, TOP-1, TOP-2 and double TOP-1/TOP-2 depletions are plotted across X chromosome genes belonging to the two categories. 1 Kb upstream of the TSS and downstream of the TES are included (Kruesi et al 2013). In the absence of TOP-2, DPY-27 enrichment is higher at genes located near strong *rex* sites. D) Metaplot of unity-normalized DPY-27 ChIP-seq signal near strong *rex* sites. 100bp bin size was used. A sliding window of 100kb and step size of 10kb was applied. The main line indicates the average and the range indicates 95% confidence interval of each window.

Closer inspection of the ChIP-seq data showed that upon TOP-2 depletion, DPY-27 peaks broadened around recruitment sites and nearby bound regions (**Figure 5B**), whereas enrichment at promoters located far away from *rex* sites was reduced (**Figure 5C**).

To quantify the progressive decrease of spreading upon TOP-2 depletion, we plotted DPY-27 ChIP-seq scores as a function of distance from the closest strong *rex* site using a sliding window (**Figure 5D**). While binding in no-tag auxin control larvae is evenly distributed, upon TOP-2 depletion, DPY-27 ChIP-seq signal progressively declines with increased distance from the *rex* sites (**Figure 5D**). This binding pattern is consistent with decreased spreading of condensin DC complexes loaded at *rex* sites and along with the Hi-C data presented above, supports the conclusion that in the absence of TOP-2, condensin DC is less processive.

### TOP-1 degradation results in accumulation of condensin DC within gene bodies

Next, we wondered how TOP-1 might contribute to condensin DC-mediated X chromosome repression (**Figure 3C**). Single-strand cleavage activity of TOP-1 is only able to remove supercoiling, thus the absence of TOP-1 should not result in the accumulation of catenations and knots (Pommier et al., 2016). To determine how TOP-1 depletion affected condensin DC binding, we performed ChIP-seq analysis of DPY-27 after one hour of TOP-1 depletion. Upon TOP-1 depletion, DPY-27 signal was evenly distributed throughout the entire chromosome regardless of the distance from a *rex* site, similar to control conditions (**Figure 5A and 5D**). Furthermore, the double degradation of TOP-1 and TOP-2 did not enhance the accumulation of condensin DC around strong *rex* sites that was observed in the single degradation of TOP-2 (**Figure 5D**). Since TOP-2, but not TOP-1 depletion affected long-range condensin DC spreading along the X chromosome, we reasoned that topological obstacles such as catenations and knots resolved by TOP-2, specifically reduce condensin DC processivity.

While long-range spreading of condensin DC is not affected by depletion of TOP-1, we noticed that DPY-27 ChIP-seq signal within gene-bodies and around transcription end sites increased specifically upon TOP-1 knockdown (**Figure 6A**). To test if increased transcription-induced supercoiling due to TOP-1 depletion causes condensin DC to accumulate in gene bodies, we first divided genes into different groups based on their expression levels measured by GRO-seq (Kruesi et al., 2013) and plotted average DPY-27 ChIP enrichment across gene bodies. Consistent with transcription driving condensin DC accumulation upon TOP-1 depletion, DPY-27 ChIP signal within the gene body was higher for genes with higher expression (**Figure 6B**). Also in support of transcription-associated supercoiling having a role in condensin DC accumulation, DPY-27 ChIP-seq enrichment within gene bodies increased with gene length, which correlates with stronger accumulation of supercoiling due to RNA Polymerase II elongating over longer distances (Liu and Wang, 1987) (**Figure 6C**).

**Figure 6.**
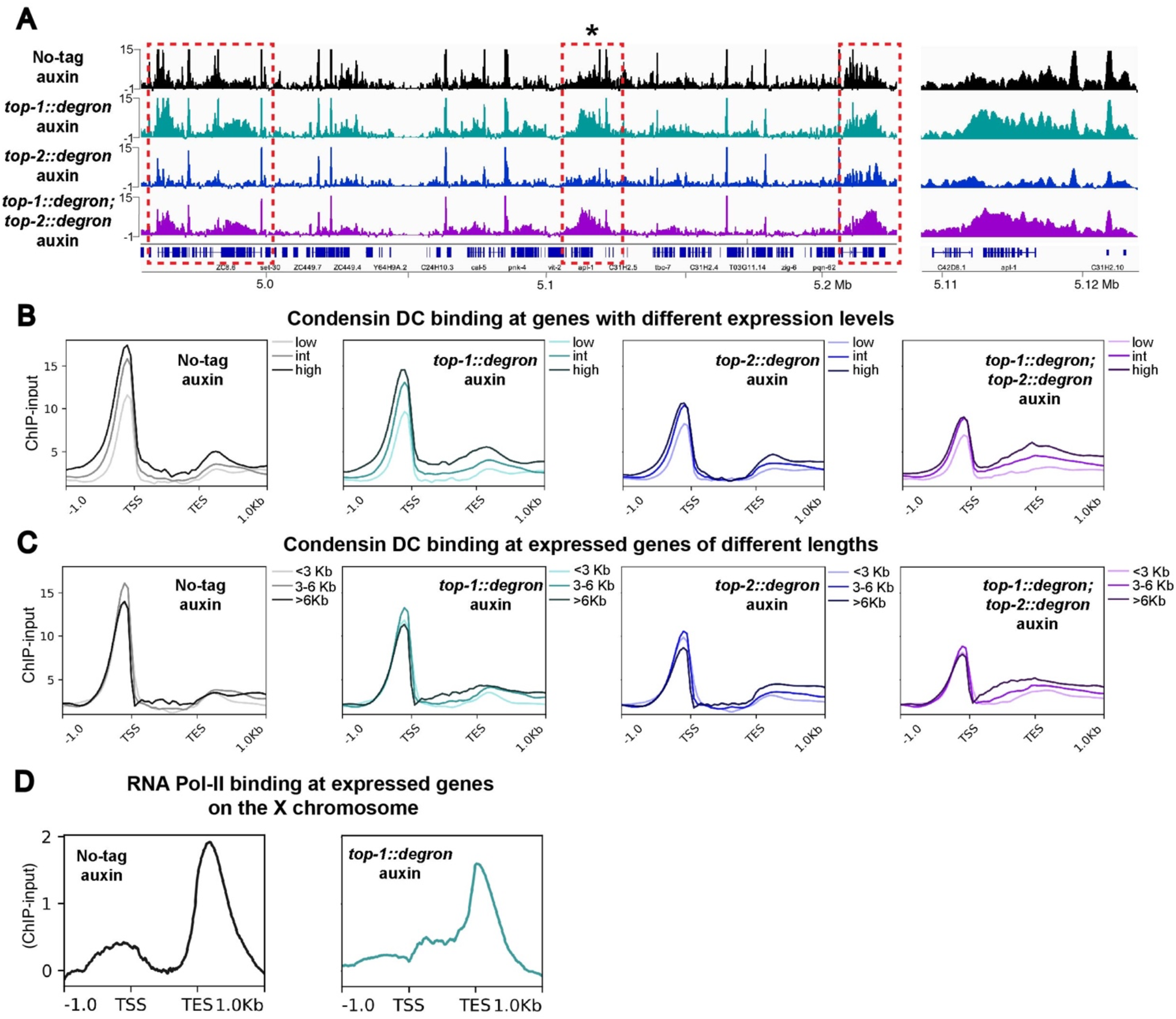
TOP-1 degradation results in accumulation of condensin DC within gene bodies. A) Left: DPY-27 ChIP-seq profile in control (no-tag auxin), TOP-1, TOP-2 and double TOP-1/TOP-2 depletions at a representative region of chromosome X. Upon TOP-1 depletion, DPY-27 signal increases within gene bodies (dashed red boxes). Right: Zoomed-in view of region indicated with an asterisk on the left. Normalized ChIP-input coverages are shown. B) Average DPY-27 ChIP-seq scores in control (no-tag auxin), TOP-1, TOP-2 and double TOP-1/TOP-2 depletion conditions are plotted across X genes grouped based on their expression level. Genes on the X chromosome were classified according to their expression level (gene-body GRO-seq signal) into three categories: low, intermediate and high. Gene-body accumulation of DPY-27 in the absence of TOP-1 increases with gene expression. C) Average DPY-27 ChIP-seq scores in control (no-tag auxin), TOP-1, TOP-2 and double TOP-1/TOP-2 depletion conditions are plotted across expressed genes on the X grouped based on their length into three categories: <3 Kb, 3-6 Kb and >6 Kb. Gene-body accumulation of DPY-27 in the absence of TOP-1 increases with gene length. D) Average RNA Pol-II ChIP-seq scores in control (no-tag auxin), and TOP-1 depletion conditions across genes on the X chromosome. TOP-1 degradation results in reduced RNA Pol-II binding at promoters and increased binding within gene bodies.

We predicted that if TOP-1 depletion results in the accumulation of transcription-induced supercoiling, it should also hinder progression of RNA Pol II (Teves and Henikoff, 2014). Indeed, RNA Pol II ChIP-seq after one hour of TOP-1 depletion showed reduced signal around transcription start sites and increased signal within gene bodies on the X and autosomes, consistent with supercoiling hindering RNA Pol II elongation (**Figure 6D and S6A**). Together these results suggest that transcription-generated supercoiling resolved by TOP-1 contributes to condensin DC translocation across gene bodies and is required for transcriptional repression.

### Distinct effects of TOP-1 and TOP-2 in condensin translocation across the X chromosomes

Both topoisomerases I and II are involved in the relaxation of supercoiling that originates from the movement of RNA Pol II along the DNA during transcription (Pommier et al., 2016). In this context, topoisomerase I and II have been shown to act in a redundant manner, although exceptions have been described for genes with high relaxation requirements such as long and highly expressed genes (Durand-Dubief et al., 2010; Joshi et al., 2012; King et al., 2013; Kouzine et al., 2013; Sperling et al., 2011). While TOP-2 depletion on its own did not result in accumulation of condensin DC at the gene bodies, depletion of both TOP-1 and TOP-2 exacerbated accumulation of condensin DC between TSS-TES (**Figure 6B and 6C**). Thus, topoisomerase II also contributes, but topoisomerase I has the major role in the resolution of transcription-induced supercoiling. Importantly, the distinct effects of TOP-1 and TOP-2 depletion on chromosome-wide distribution of condensin DC suggests that long-range translocation, which is sensitive to TOP-2, is not affected by gene-body accumulation of condensin DC upon TOP-1 depletion. In other words, condensin DC translocation across transcribed genes was hindered in the absence of TOP-1, but this blockage did not affect spreading from recruitment sites (**Figure 5A and 5D**). Furthermore, stronger gene-body accumulation of condensin DC in the double TOP-1;TOP-2 depletion did not result in a further reduction of spreading from recruitment sites, like one would expect if long-range spreading required the complex to translocate across transcribed genes (**Figure 5D**). This suggests that condensin DC complexes spreading across long distances from recruitment sites can bypass transcription-induced supercoiling whereas condensin DC complexes translocating within gene bodies are susceptible to such topological constraints.

## Discussion

In this study we provide *in vivo* evidence for a role of topoisomerases in the regulation of condensin DC translocation during interphase. Our data indicate that topoisomerases I and II regulate condensin DC translocation at different genomic scales. TOP-1 is required for the local translocation of condensin DC through genes which necessitates the resolution of supercoiling primarily generated in the context of transcription. TOP-2 is required for long-range translocation of condensin DC across hundreds of kb regions along the X chromosomes, likely through the removal of DNA catenates and knots. These observations provide important insights into several fundamental questions in the field of chromosome structure and function.

### TOP-2 promotes condensin DC long-range translocation and loop formation

Topoisomerase II and condensin are key players in the resolution, compaction and segregation of chromosomes during cell division. TOP-2 and condensin subunits have been identified as the most abundant non-histone proteins on mitotic chromosomes (Maeshima and Laemmli, 2003) and are part of the minimal components required to make mitotic-like chromosome structures *in vitro* (Shintomi et al., 2015). Modeling experiments proposed that as SMC complexes extrude, they could push and localize catenations and knots, thus promoting their relaxation by TOP-2 (Orlandini et al., 2019; Racko et al., 2018). One simulation study showed that reducing DNA chain passing with fixed loop extrusion parameters enhances TADs, thus it was proposed that topological constraints reinforce the impact of loop extrusion (Nuebler et al., 2018). However, our results indicate that topological constraints arising specifically upon TOP-2 depletion can in turn affect condensin-mediated DNA loop extrusion. Future simulations should consider the possibility that the properties of loop extrusion depend on the frequency of chain passing in order to understand the contribution of DNA topology and loop extrusion on the 3D genome folding features.

Although we interpret our results using DNA loop extrusion as a framework, it is possible that long-range condensin DC translocation along chromatin *in vivo* uses other mechanisms such as diffusion. Since condensin DC spreads linearly over Mb distances in X;autosome fusion chromosomes (Ercan et al., 2009), this non-extrusion mechanism should also be linear. In this case, in the absence of TOP-2, X chromosome structure should change in a way that affects translocation, concentrating condensin DC around the strong recruitment sites to produce the ChIP-seq profiles we observe (**Figure 5A**).

### Continuous addition and removal of DNA knots by TOP-2 may regulate condensin processivity

*In vivo* there seems to be a limit in the capacity of condensins to push through intertwined chromatin. The extent of this limitation is unclear but one hour of TOP-2 depletion led to an overall decrease in the distance-range of DNA contacts on the X chromosome and strongly reduced condensin DC spreading. Thus, we propose that *in vivo* loop extrusion by condensins continually requires TOP-2 strand passage activity. The requirement of TOP-2 for condensin DC translocation in larvae suggests that topological constraints are generated even in somatic cells that have largely completed their divisions. One possibility is that TOP-2 constantly produces short-lived entanglements. Positive supercoiling resulting from transcription was shown to increase the formation of DNA knots by TOP-2, and it was suggested that after positive supercoiling is relaxed, these knots would get resolved by TOP-2 making them short-lived (Valdés et al., 2019). The genome-wide frequency and distribution of such transcription-induced knots is not known, but it is possible that TOP-2 acute depletion results in unresolved knots that have the potential to halt condensin translocation.

### The strong *rex* sites act as TAD boundaries and recruit TOP-2

Several studies support a model in which condensin activity confers directionality to the strand passage reaction performed by TOP-2, favoring decatenation over catenation (Baxter et al., 2011; Charbin et al., 2014; Dyson et al., 2020; Piskadlo et al., 2017; Sen et al., 2016), thus contributing to chromatin unknotting. Modeling experiments proposed that the activity of condensin leads to the recruitment of TOP-2 to highly entangled regions (Orlandini et al., 2019). Our results support this hypothesis by showing that in the absence of condensin DC, the binding of TOP-2 to strong *rex* sites that also serve as the TAD boundaries on the X chromosome is reduced. In addition, the ectopic insertion of a recruitment element that establishes a new TAD boundary (Jimenez et al., 2021), is sufficient to recruit TOP-2. Therefore, it is possible that condensin DC induces topological configurations that recruit TOP-2 to the strong *rex* sites. Whether TOP-2 recruitment to TAD boundaries underlies its enrichment on X chromosome promoters remains unclear. It is possible that TOP-2 is pushed out of the recruitment sites and spreads linearly with condensin DC along the X chromosomes. It is also possible that active and continuous recruitment of TOP-2 to the *rex* sites could contribute to concentrating TOP-2 to the X chromosome territory in the nucleus. Alternatively, the action of condensin DC at X chromosomal promoters could change the topology of chromatin into a configuration that is recognized by TOP-2.

### TOP-1 is required for condensin DC translocation across transcribed gene bodies

The relaxation of transcription-induced supercoiling by topoisomerases is required for transcription elongation (Durand-Dubief et al., 2010; Pommier et al., 2016). Here we show that in *C. elegans*, TOP-1 is the main relaxer of transcription-generated supercoiling across the genome and propose that on the X, this relaxation activity is important for the translocation of condensin DC across transcribed genes. Transcription has been proposed to affect SMC translocation in different ways. On one hand, high concentrations of transcribing RNA Pol II have been proposed to act as permeable barriers that slow condensin loop extrusion (Brandão et al., 2019; Tran et al., 2017). On the other hand, transcription has been proposed to be a driving force for cohesin translocation, either by RNA Pol II directly pushing the cohesin ring or through the energy originated from transcription-induced supercoiling (Busslinger et al., 2017; Davidson et al., 2016; Rusková and Račko, 2021). Therefore, condensin DC gene body accumulation upon TOP-1 depletion could be the result of stalled RNA Pol II complexes not pushing condensin DC or blocking its translocation.

### TOP-1 and TOP-2 contribute to condensin-DC mediated transcription repression

The fact that disruption of both TOP-1 and TOP-2 caused X chromosome derepression suggests that condensin DC mediated chromosome-wide repression requires both long-range linear spreading and local binding to genes. Long-range translocation from recruitment sites through loop extrusion allows spreading of the complex throughout the X chromosome, creating prominent TAD structures (Crane et al., 2015). The level of condensin DC spreading within large genomic windows correlates with the level of repression in X-autosome fusion chromosomes, demonstrating the link between binding and repression (Street et al., 2019). Thus TOP-2 depletion, which antagonizes spreading, would interfere with repression. At the gene level, gene expression analysis in dosage compensation mutants showed no strict correlation between the level of repression and condensin DC binding at promoters, invoking a repression mechanism acting at a distance (Jans et al., 2009). While better experimental approaches to measure direct effect of condensin DC binding are required, it remains possible that local translocation of condensin DC could interfere with the transcriptional machinery to reduce RNA Pol II recruitment at X chromosomal promoters (Kruesi et al., 2013). This mechanism may be related to transcription-generated supercoiling of genes, as revealed by accumulation of condensin DC upon TOP-1 depletion.

### The distinct effects of TOP-1 and TOP-2 depletion on condensin DC distribution support distinct modes of condensin DC translocation over long and short distances

Strikingly, the effect of condensin DC accumulation at gene bodies upon TOP-1 depletion was local and did not reduce its long-range linear spreading from the recruitment sites. Thus, it is likely that condensin DC has an additional mode of association with chromatin that happens locally and is susceptible to topological constraints such as transcription-induced supercoiling. Contrary to this local gene body association, spreading of condensin from *rex* sites do not seem to be sensitive to topological constraints arising in the absence of TOP-1. What determines the different behaviors of condensin DC? One distinguishing factor between condensin DC complexes found at recruitment sites and complexes found at gene promoters is the presence of the main condensin DC recruiter protein, SDC-2, which shows high binding at *rex* sites and limited spreading across the X chromosome (Ercan et al., 2009). Similar to the cohesin loader, NIPBL, SDC-2 could associate with extruding condensin DC complexes and act as a processivity factor allowing condensin to overcome obstacles. Condensin DC extruding complexes that dissociate from SDC-2, could engage in a different mode of DNA translocation that involves movement through the gene in a transcription-coupled manner, and is sensitive to DNA supercoiling.

### A model for DNA catenation and transcription-induced supercoiling regulating *in vivo* condensin translocation

Here we present a model in which condensin DC’s linear spreading over long distances requires resolution of DNA entanglements by TOP-2 and translocation over genes requires the resolution of DNA supercoiling by TOP-1 (**Figure 7**). In wild type cells, catenations may be resolved by TOP-2 at a high rate, but as condensin continues to move along chromatin, chances of encountering additional blocks would increase, resulting in reduced binding with increased distance from the *rex* sites. This would explain the gradual decrease in condensin DC spreading into the autosomal region of X;A fusion chromosomes over 1-3 Mb distances (Ercan et al., 2009). SDC-2 and SDC-3 proteins bound at strong *rex* sites could support condensin DC translocation over Mb-scale distances. Even distribution of *rex* sites, ∼1 Mb apart across the X, would accomplish the chromosome-wide distribution of condensin DC. At the local scale, another pool of condensin DC could be translocating over genes in a manner sensitive to transcription-induced supercoiling and thus require TOP-1. In yeast, condensin was reported to bind active gene bodies during mitosis to prevent transcription induced chromosome segregation defects (Sutani et al., 2015). Mechanism of transcription repression by condensin DC may be related to this second translocation activity and co-opted from a mitotic function by condensins.

**Figure 7.**
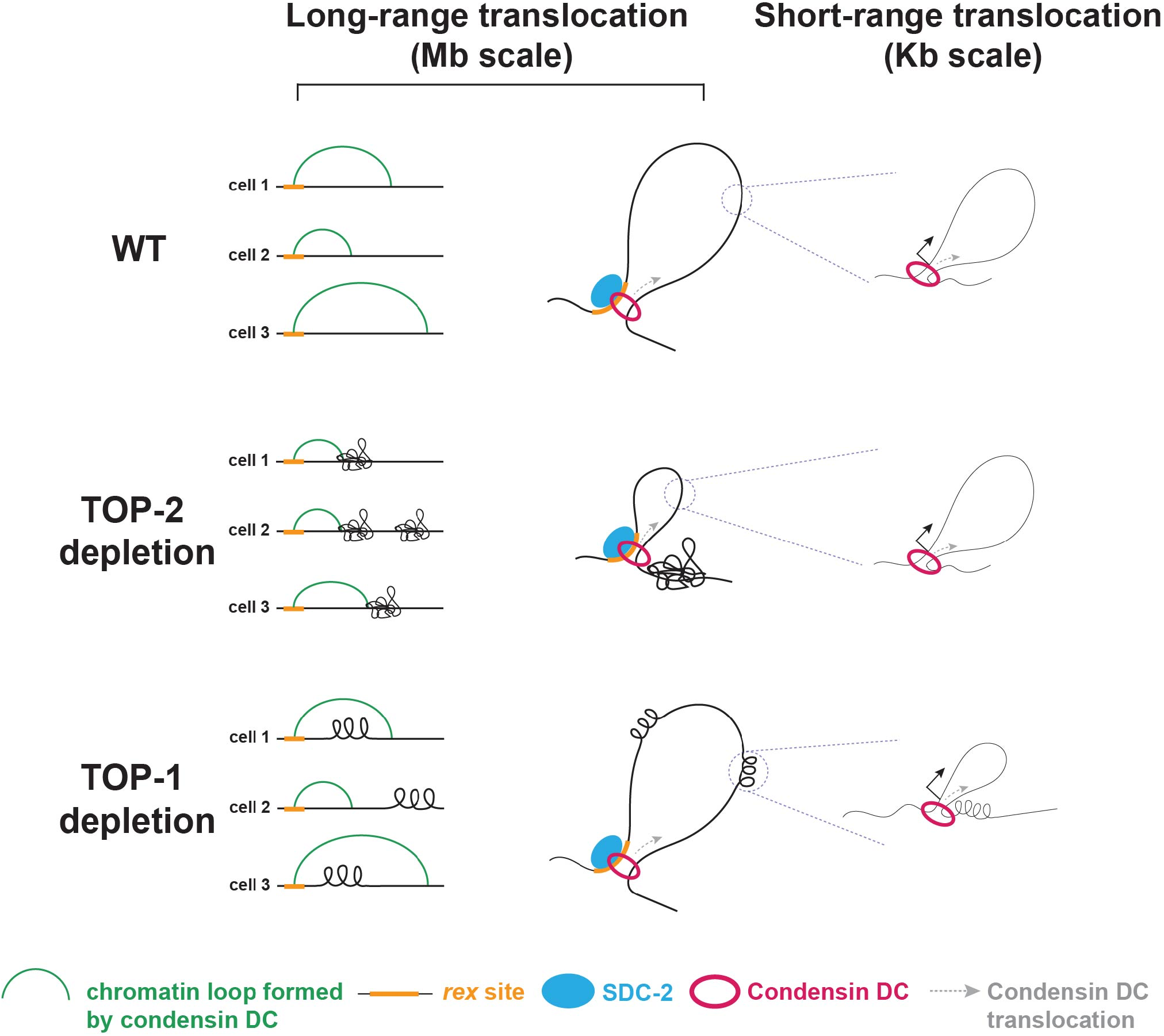
Model for regulation of condensin DC translocation by topoisomerases I and II. At *rex* sites SDC-2 may promote condensin DC processivity enabling it to spread over Mb-scale distances. This long-range translocation of the complex requires the resolution of topological constraints such as DNA catenates and knots by TOP-2 but is capable of bypassing transcription-induced supercoiling that accumulates in the absence of TOP-1. Some condensin DC complexes could dissociate from SDC-2 at promoters and engage in a shorter scale translocation within gene-bodies that cannot overcome transcription-induced supercoiling.

Our model opens a new line of research to understand the mechanism by which condensin and topoisomerase II function together. While the role of condensin regulating topoisomerase II is better understood, here we show that topoisomerase II in turn, promotes condensin processivity. Thus upon topoisomerase II depletion or inhibition, blocked condensin complexes may contribute to previously reported condensation defects (Piskadlo and Oliveira, 2017). Our work also contrasts condensin to cohesin, another SMC complex with loop extrusion activity (Golfier et al., 2020). A recent study has shown that drug inhibition of both topoisomerase I and II activity may reduce cohesin processivity (Neguembor et al., 2021). It is possible that unlike condensin which has two distinct modes of translocation including one that “skip over” genes, cohesin may have to move through transcribed regions and is thus sensitive to DNA supercoiling. In summary, our study sheds light into how different types of topological constraints impact condensin translocation and function. Deciphering how loop extrusion by SMCs is impacted by factors inherent to the chromatin fiber will contribute to the better understanding of the mechanisms by which the genome is dynamically organized throughout the cell cycle.

## Acknowledgements

AKM, SE, DO and research in this manuscript were supported by NIGMS of the National Institutes of Health under award numbers R01 GM107293 and R35 GM130311. Jun Kim was partially supported by NIGMS Predoctoral Fellowship T32HD007520. We thank Paul Maddox for the TOP-2 antibody and the *top-2::sfGFP* strain. We thank the gencore at the NYU Center for Genomics and Systems Biology for sequencing and raw data processing.

## Author Contributions

AKM and SE conceptualized the project and designed experiments. AKM, JK and SS performed experiments. AKM, JK and DO performed bioinformatics analysis. AKM and SE wrote the manuscript.

## Declaration of interests

The authors declare no competing interest.

## Materials and methods

### Worm strains and growth

Strains, primers, sgRNAs, and Sanger sequencing results are provided in the supplemental tables. Worms were grown and maintained at 20-22°C on NGM plates containing *E. coli* strains OP50-1 and/or HB101. Mixed developmental stage embryos were obtained by bleaching gravid adults. To isolate synchronized L2/L3 worms, gravid adults were bleached and embryos were hatched overnight in M9 buffer. The resulting starved L1s were grown for 24 hours at 22°C. For *glp-1* adult worms, gravid adults grown at the permissive temperature (15°C) were bleached and starved L1s were grown at the restrictive temperature (25°C) for three days.

Auxin treatment: synchronized L2/L3 worms were washed three times with M9 and split into two aliquots. Half of the worms were transferred to NGM plates containing 1mM of indole-3-acetic-acid (Fisher 87-51-4) prepared as in (Zhang et al., 2015). The other half were placed in normal NGM plates (no-auxin control) for the indicated time. Worms were then washed one time with M9 and processed accordingly to future application. For ChIP and Hi-C, L3s were incubated in 2% formaldehyde for 30 minutes, followed by quenching with glycine, one wash with M9 and two washes with PBS, PMSF and protease inhibitors. For RNA-seq, worms were stored in Trizol.

#### Generation of *top-1::degron::GFP*

The degron-GFP tag was inserted at the 3’ end of *top-1* (M01E5.5) using the CRISPR/Cas9 system. A 20 bp crRNA (AM56) was designed to target the end of the last *top-1* exon. Repair templates consisting of a 15 bp flexible linker (GlyGlyGlyGlySer) and the degron-GFP tag flanked by 120 bp single stranded overhangs complementary to *top-1* were generated as previously described (Dokshin et al., 2018) by PCR using AM57F&R and AM04&AM20 primers and pLZ29 as a template. The injection mix containing *S*.*pyogenes* Cas9 3NLS (10 mg/ml, IDT), crRNA (2 nmol, IDT), tracrRNA (5 nmol, IDT), dsDNA donors, and pCFJ90 (pharynx mCherry marker) was prepared as described in (Dokshin et al., 2018). Around 30 F1s that were positive for the co-injection marker were transferred to individual plates. F2 worms were screened by PCR using primers AM58F&R. PCR products of the expected size were sequenced to confirm in-frame insertion. Sanger sequencing results are provided in Supplemental File 1.

#### Generation of *top-2::degron::GFP*

Same as for *top-1::degron::GFP* with the following reagents:

crRNA: AM30

Primers to generate repair template: AM22F&R

Genotyping primers: SS02F&R

#### Generation of *dpy-27::degron::GFP*

Same as for *top-1::degron::GFP* with the following reagents:

crRNA: LS37

Primers to generate repair template: AM21F&R

Genotyping primers: LS40F&R

### Genomic data access

Genomic data is available at Gene Expression Omnibus (GEO) series numbers GSE188851. Accession numbers of the data sets generated in this study are listed in Supplemental table 4,5 and 6.

### ChIP-seq

Two biological replicates with matching input samples as reference were performed for each experiment (Supplemental file 1). Embryos, L2/L3s and adult worms were dounce-homogenized in FA buffer (50 mM HEPES/KOH pH 7.5, 1 mM EDTA, 1% Triton X-100, 0.1% sodium deoxycholate; 150 mM NaCl) and sonicated in 0.1% sarkosyl to obtain chromatin fragments between 200 and 800 bp. Protein concentration was determined using Bradford assay (Biorad 500-0006). As in (Ercan et al., 2007), 2 mg of protein extract and 3 to 10 ug of antibody was used per ChIP. Antibodies used in this study are listed in Supplemental file 1. For library preparation, half of the ChIP DNA and 30 ng of input DNA were ligated to Illumina TruSeq adapters and amplified by PCR. Library DNA between 250 and 600 bp was gel purified. Single-end 75 bp sequencing was performed using the Illumina NextSeq 500 at the New York University Center for Genomics and Systems Biology, New York, NY.

#### ChIP-seq data processing and analysis

Bowtie2 version 2.4.2 was used to align 75 bp single-end reads to WS220 with default parameters (Langmead and Salzberg, 2012). Bam sorting and indexing was performed using samtools version 1.11 (Danecek et al., 2021). BamCompare tool in Deeptools version 3.5.0 was used to normalize for the sequencing depth using CPM and create ChIP-Input coverage and ChIP/Input ratios with a bin size of 10 bp and 200 bp read extension (Ramírez et al., 2016). Only reads with a minimum mapping quality of 20 were used, and mitochondrial DNA, PCR duplicates, and blacklisted regions were removed (Amemiya et al., 2019). The average coverage data was generated by averaging ChIP-Input enrichment scores per 10 bp bins across the genome. Heatmaps and average-profile plots of ChIP-seq scores across different annotations were produced using Deeptools in Galaxy (doi:10.1093/nar/gkw343). Average score over the regions was calculated using 50 bp non-overlapping bins. In figure 5C, 6B-C-D and S6, genes were scaled to 1000 bp.

For alignments and sliding window analysis: Input subtracted ChIP signal across X-chromosome is binned into 100 bp resolution. The signal across the X-chromosome is normalized to unity. Each bin on X-chromosome is assigned to the closest strong *rex* site defined in (Albritton et al., 2017). Then, the normalized signal is plotted as a function of linear distance away from rex sites with sliding window size of 100 kb and step size of 10 kb, where the main line indicates the average and the wider range indicates the 95% confidence interval of the window.

For motif analysis: MEME (version 5.3.3) (Machanick and Bailey, 2011) was used to find DNA sequence motifs enriched at TOP-2 binding sites. 150 bp sequences centered on TOP-2 ChIP summits were analyzed using the parameters: minimum motif size 6 bp; maximum motif size 12 bp; expected motif occurrence set to any number of occurrences and other parameters set to default. The position weight matrix (pwm) of the top enriched motif was saved from the MEME output and used in PWMScan (Ambrosini et al., 2018) to scan the *C. elegans* ce10 genome for genome wide occurrences of the motif. The p-value cut-off was set to 1e-5, and other parameters were set as default. The hits obtained with PWMScan were used in R to get the sequence motif chromosomal distribution.

For GC content analysis: The GC content of 250-bp sequences centered around the TSS was obtained using bedtools (version 2.27.1) NucBed function with the *C. elegans* ce10 genome and default parameters. The TOP-2 ChIP binding for each region was obtained using deeptools (version 3.3.1) multiBigWigSummary function with the ‘BED-file’ option and other parameters as default.

### mRNA-seq

L3 larvae were collected for two biological replicates per condition and stored in Trizol (Invitrogen) at −70°C. Total RNA was purified following manufacturer’s instructions after freeze-cracking samples five times. RNA was cleaned up using Qiagen RNeasy MinElute Cleanup kit and quantified using Nanodrop. 1 µg of RNA was run on 1% agarose gel complemented with bleach to assess quality (Aranda et al., 2012). mRNAs were purified using Sera-Mag Oligo (dT) beads (Thermo Scientific) from 10 µg of total RNA. Stranded Illumina libraries were prepared as previously described (Kramer et al., 2015). Single-end 75 bp sequencing was performed using the Illumina NextSeq 500 at the New York University Center for Genomics and Systems Biology, New York, NY.

#### RNA-seq data processing and analysis

We aligned reads to the WS220 genome version using HISAT2 version 2.2.1 (Kim et al., 2019) with the parameter --rna-strandness R. Count data was calculated using HTSeq version 0.13.5 (Anders et al., 2015). The raw counts were normalized FPKM using cufflinks version 2.2.1 (Roberts et al., 2011), and then FPKM was converted to TPM. The raw counts were used for the R package DESeq2 version 1.30.0 (Love et al., 2014). Violin and box plots were produced in R using ggplot2 (http://ggplot2.org). Outliers were not plotted.

### Hi-C

Crosslinked worms collected as described above were resuspended in 20 ul of PBS per 50ul of worm pellet then dripped into liquid nitrogen containing mortar. The worms were grounded with pestle until fine powder. Grounded worms were crosslinked again in 2% formaldehyde using the TC buffer as described by the Arima Hi-C kit, which uses two 4-base cutters, DpnII and HinfI. The Arima manufacturer’s protocol was followed including the recommended method of the library preparation using KAPA Hyper Prep Kit. Paired-end 150 bp sequencing was performed using the Illumina Novaseq 6000 at the New York University Center for Genomics and Systems Biology, New York, NY.

#### HiC data processing and analysis

##### Hi-C data analysis

The Hi-C data was mapped to ce10 (WS220) reference genome using default parameters of the Juicer pipeline version 1.5.7 (Durand et al., 2016). The biological replicates were combined using juicer’s mega.sh script. The mapping statistics from the inter_30.txt output file are provided in Supplementary Table 1. The inter_30.hic outputs were converted to h5 using the hicConvertFormat of HiCExplorer version 3.5.1 (Ramírez et al., 2018; Wolff et al., 2020, 2018) for sample-to-sample depth normalization and genome-wide matrix balancing. The inter_30.hic files were first converted to cool files, and the correction method was removed using the -- correction_name none option. Then, cool files were converted to h5 files to be used in HiCExplorer. The count values of each replicate were normalized to match those of the shallowest matrix using hicNormalize with the option --smallest. The same method was used for the combined matrix. Lastly, the hicCorrectMatrix function was applied to each matrix to correct for visibility bias of genomic bins with the following parameters: --correction_method ICE, -t 1.7 5, --skipDiagonal, -- chromosomes I II III IV V X. The distance decay curves were generated using the 500 bp-binned balanced matrix using hicPlotDistVsCounts with parameters --perchr, maxdepth 20,000,000. The outputs from --outFileData were plotted in R. To analyze X-specific changes, we calculated P(s,chrX)/P(s,chrA) by dividing the P(s) of the X chromosome by the average P(s) of all autosomes at every distance. The pile-up analysis used the 17 strong *rex* sites defined in (Albritton et al., 2017). For off-diagonal *rex-rex* metaplot, the 10 kb-binned matrix with the hicAggregateContacts function of HiCexplorer was used with parameters: --range 100000:3000000, --avgType mean, -- transform obs/exp, --plotType 3d, --vMin 0.8 --vMax 2. For on-diagonal rex metaplot, the 500bp-binned balanced matrix was converted to cool format and computed using cooltools version 0.4.0 (https://github.com/open2c/cooltools).

### Microscopy

For direct comparison, *top-1::degron::GFP* and *top-2::degron::GFP* worms incubated with and without auxin were aligned side by side prior to imaging as previously described in (Zhang et al., 2015) (**Figure 3B**). For *glp-1* mutants (**Figure S1B**), adult worms grown at the permissive (16°C) and restrictive (25°C) temperatures were immobilized with 100 mM sodium azide immediately before imaging. Images were acquired with a Zeiss Axio Imager A2 microscope using a 10x and 20x objective and the AxioVision Rel.4.8 software.

## Notes

### Competing Interest Statement

The authors have declared no competing interest.

