## Supplemental Information for "Topoisomerases I and II facilitate condensin DC translocation to organize and repress X chromosomes in *C. elegans*"

### Supplementary figure 1

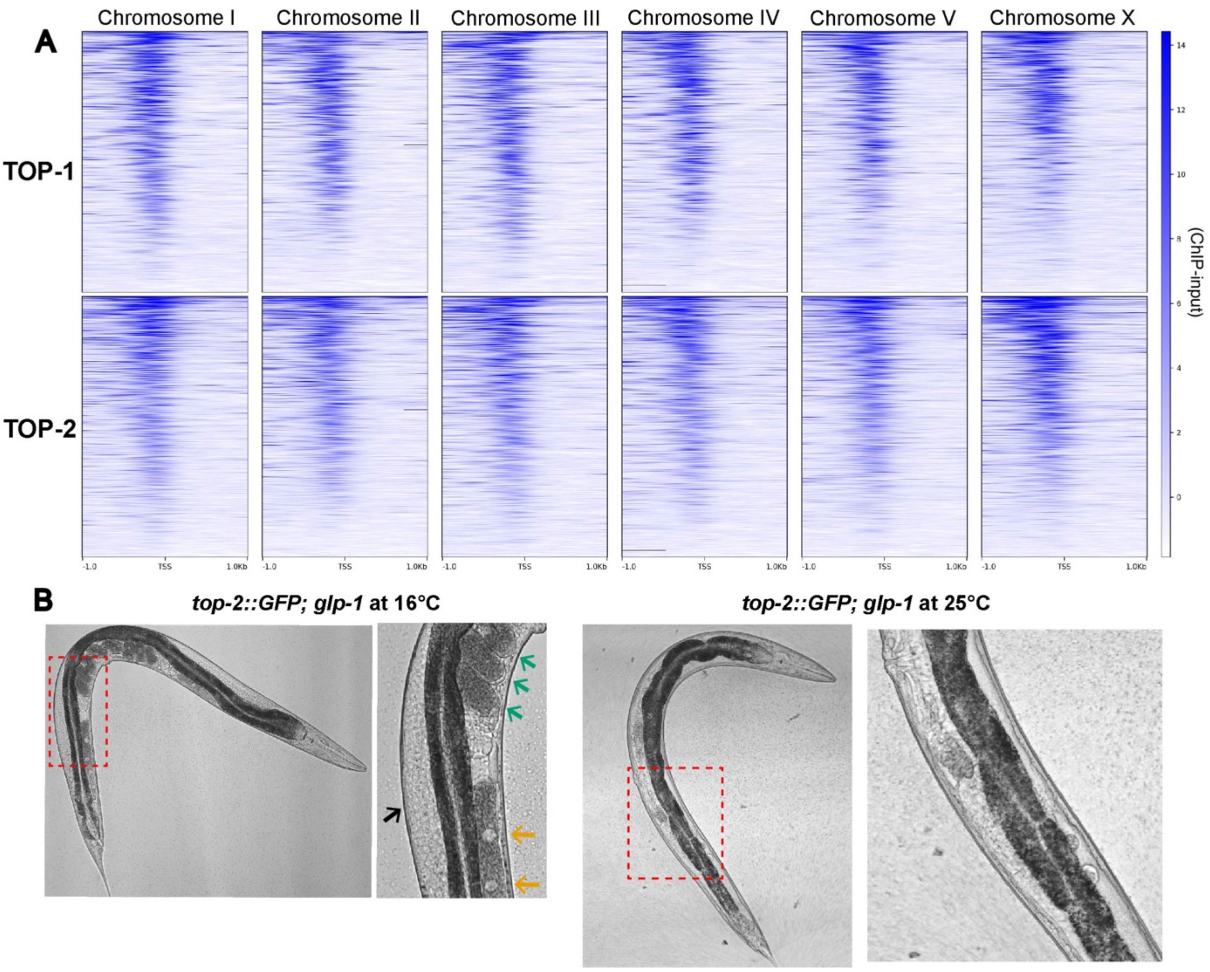


**Figure S1.**

A) Heatmap showing TOP-1 and TOP-2 ChIP-seq signals across TSSs on all chromosomes (Kruesi et al., 2013).

B) Images of *top-2::sfGFP; glp-1(q224)* adults that were grown at the permissive (16°C) and restrictive (25°C) temperatures. *glp-1(q224)* worms lack a germline when grown at the restrictive temperature. Black arrow indicates the germline (surface view). Orange arrows indicate oocytes. Green arrows indicate fertilized embryos. All these structures are missing in worms grown at 25°C.

### Supplementary Figure 2

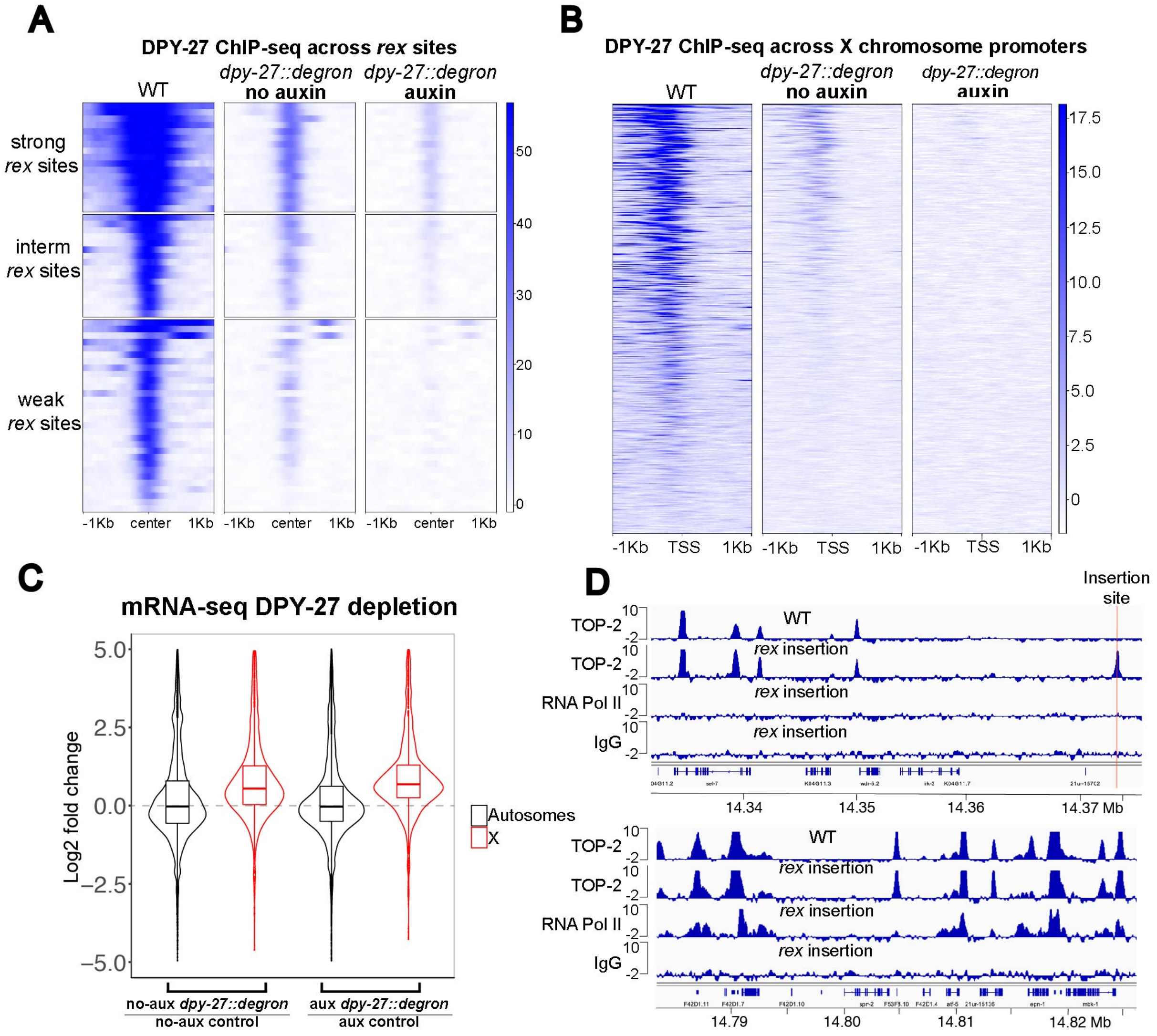


**Figure S2.**

A) Heatmap showing DPY-27 ChIP-seq signal across a 2 Kb window centered around 64 *rex* sites previously described (Albritton et al., 2017) in WT, *dpy-27::degron::GFP* worms without auxin and *dpy-27::degron::GFP* worms treated with auxin for 60min. Insertion of the degron-GFP tag in the presence of the TIR-1 protein impaired DPY-27 function and reduced binding to the X chromosome without the addition of auxin.

B) Heatmap showing DPY-27 ChIP-seq signals across GRO-seq defined TSSs (Kruesi et al., 2013) in the WT, *dpy-27::degron::GFP* strain without auxin and *dpy-27::degron::GFP* worms treated with auxin for 60min.

C) mRNA-seq data of *dpy-27::degron::GFP* worms that were treated with or without auxin. Fold changes were calculated between no-auxin *dpy-27::degron::GFP* and no-auxin control and between 120 min auxin *dpy-27::degron::GFP* and 120 min auxin control. The distributions of log2 fold changes are shown for Autosomes and X chromosome.

D) ChIP-seq profiles of TOP-2 in WT and TOP-2, RNA Pol-II and IgG in a strain carrying an ectopic *rex-8* insertion on the X chromosome. Profiles around the insertion site (top panel) and an additional representative (bottom panel) region of the X chromosome are shown.

**Supplementary figure 3**

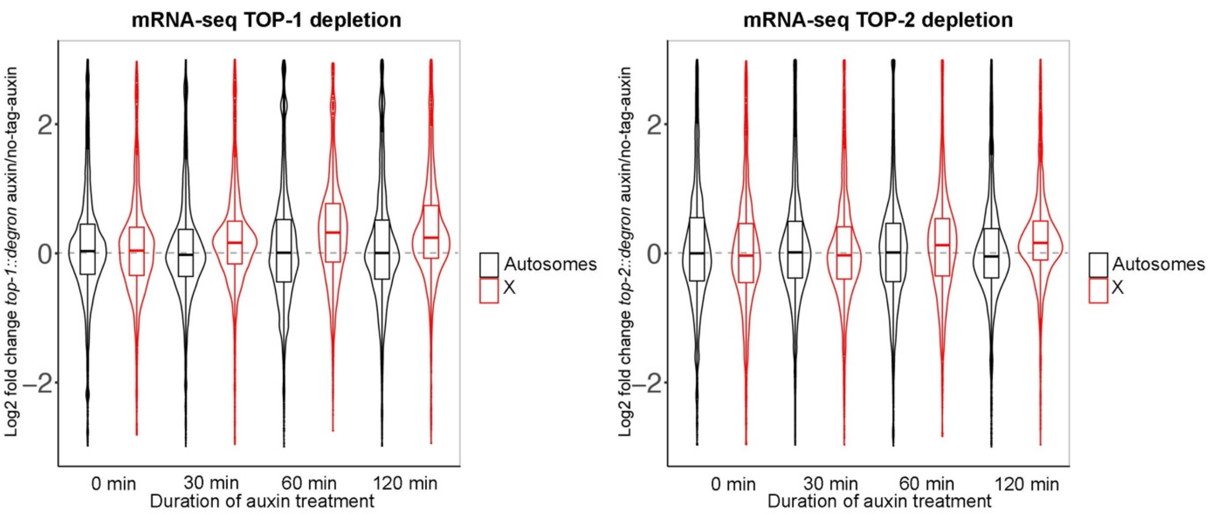


**Figure S3.**
mRNA-seq was performed after 30 min, 60 min and 120 min auxin-mediated depletion of TOP-1 and TOP-2. Distribution of log2 fold changes between the degron-auxin and no-tag-auxin conditions are shown for autosomes and X chromosomes.

### Supplementary figure 4

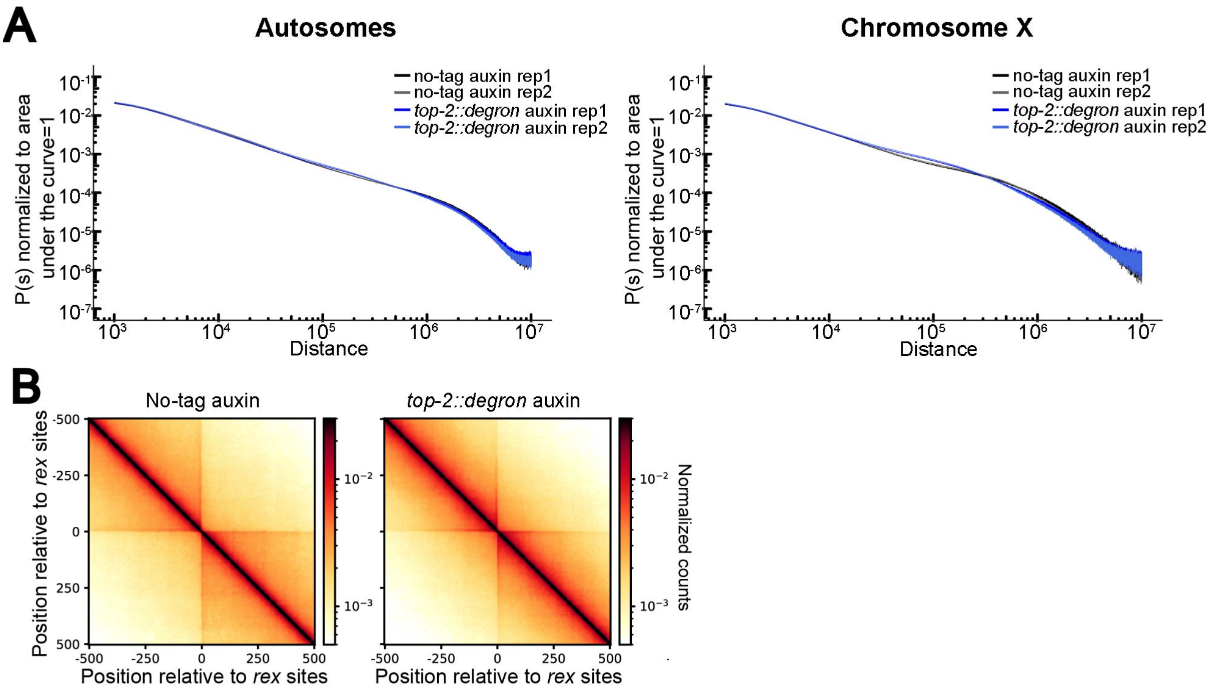


**Figure S4.**

A) Distance decay curve showing the relationship between 500bp binned genomic distance and average contact probability computed for all autosomes (left) and the X-chromosome (right) in the no-tag auxin control and *top-2::degron* auxin .

B) Pile-up analysis showing the average Hi-C map +/- 500-kb surrounding the annotated 17 strong *rex* sites.

### Supplementary figure 5

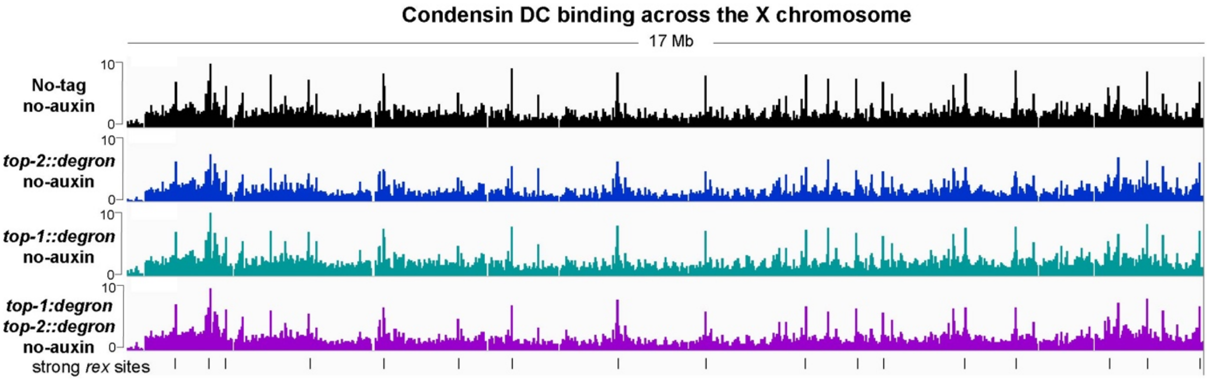


**Figure S5.**
X chromosome view of DPY-27 ChIP-seq profile in control (no-tag), *top-2::degron::GFP*, *top-1::degron::GFP* and double *top-1::degron::GFP; top-2::degron::GFP* worms that were incubated in no-auxin plates for 60 minutes. Black lines at the bottom indicate the location of strong *rex* sites.

### Supplementary figure 6

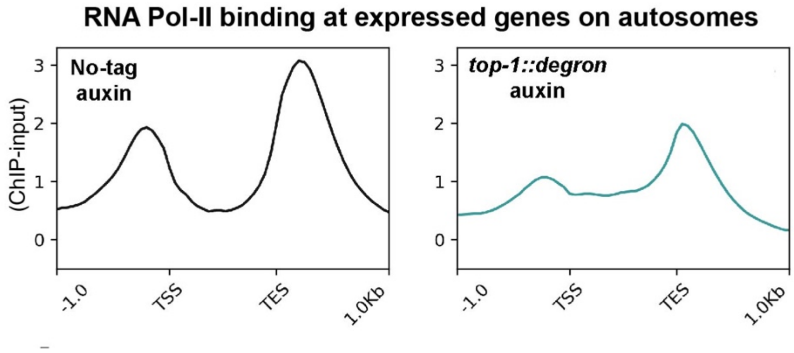


**Figure S6.**
Average RNA Pol-II ChIP-seq scores in control (no-tag auxin), and TOP-1 depletion conditions across expressed genes on the autosomes. TOP-1 degradation results in reduced RNA Pol-II binding at promoters and increased binding within gene bodies.

**Table S1. List of the *C. elegans* strains used in this study**

| *strain name* | *short name* | *short description* | *strain genotype* | *Source* |
|---|---|---|---|---|
| ERC82 | AM01 | *dpy-27::degron::GFP* endogenous location in CA1200. Dumpy phenotype | ers54[dpy-27::degron::GFP] III; ieSi57 [eft-3p::TIR1::mRuby::unc-54 3'UTR + Cbr-unc-119(+)] II | this study |
| ERC83 | SS01 | *top-2::degron::GFP* endogenous location in CA1200. Fully complementing function | ers55[top-2::degron::GFP] II; ieSi57 [eft-3p::TIR1::mRuby::unc-54 3'UTR + Cbr-unc-119(+)] II | this study |
| ERC84 | AM05 | *top-1::degron::GFP* endogenous location in CA1200. Fully complementing function | ers56[top-1::degron::GFP] I; ieSi57 [eft-3p::TIR1::mRuby::unc-54 3'UTR + Cbr-unc-119(+)] II | this study |
| ERC69 | LS05 | *ectopic rex-8* inserted in chromosome X | ersIs33[X:11093924-11094281[rex-8], X:14373128] | Jimenez et al, 2021 (preprint) |
| ERC38 | SEA03 | *rex-41* deletion | ers30[delX:17544437-17544484, delX:17545624-17545624] | Albritton et al, 2017 |
| JK1107 | | temp sensitive *glp-1* mutant, viable at 15°C | *glp-1(q224)* III | CGC |
| MDX53 | | *top-2::sfGFP* endogenous location. Fully complementing function | top-2::sfGFP-3xFLAG; mCherry::H2B | Maddox lab (Ladouceur et al., 2017) |
| CA1200 | | Single copy transgene inserted into chromosome II (oxTi179) expressing modified Arabidopsis thaliana TIR1 tagged with mRuby in the soma | ieSi57 [eft-3p::TIR1::mRuby::unc-54 3'UTR + Cbr-unc-119(+)] II | CGC |

**Table S2. List of the antibodies used in this study**

| Target | Antibody | Antibody information | Antigen | Source of antibody | Reference |
|---|---|---|---|---|---|
| DPY-27 | JL00001 | Rabbit polyclonal | 1-409 aa | Covance Research Products Inc Cat# JL00001_DPY27, RRID:AB_2616039 | Ercan et al., 2007 |
| GFP | ab290 | Rabbit polyclonal | Recombinant full-length protein corresponding to GFP. Green fluorescent protein (GFP) from Aequorea victoria. | Abcam | |
| RNA Pol-II | ab817 | mouse monoclonal | RNA polymerase II CTD repeat YSPTSPS | Abcam | |
| RNA Pol-II | 05-952-I-100UG | mouse monoclonal | RNA polymerase II CTD repeat YSPTSPS | Millipore | |
| TOP-2 | | Rabbit polyclonal | 1,350-1,470 aa *C. elegans* TOP-2 | | (Ladouceur et al., 2017) |
| IgG | ab46540 | Rabbit polyclonal | mouse IgG heavy and light chains | Abcam | |

**Table S3. List of the primers used in this study**

**Table S4. Information for ChIP-seq data generated in this study**

**Table S5. Information for RNA-seq data generated in this study**

**Table S6. Information for Hi-C data generated in this study**

**Table S7. Sanger sequencing results for strain ERC84 (*top-1::degron::GFP*)**

**Table S8. Sanger sequencing results for strain ERC83 (*top-2::degron::GFP*)**

**Table S9. Sanger sequencing results for strain ERC82 (*dpy-27::degron::GFP*)**
